# Polarity-dependent modulation of sensory circuits by cerebellar tDCS: local and distant effects

**DOI:** 10.1101/2025.05.12.653393

**Authors:** Carlos Andrés Sánchez-León, Isabel Cordones, Alba Jiménez-Díaz, Guy Cheron, Javier F Medina, Javier Márquez-Ruiz

**Author notes:** **Corresponding author:** Javier Márquez-Ruiz, Ph.D., Brain Stimulation translational Laboratory Universidad Pablo de Olavide, Ctra. de Utrera, km. 1 41013-Sevilla, Spain. **Declaration of interests**: The authors declare no competing financial interests.

## Abstract

Cerebellar transcranial direct current stimulation (Cb-tDCS) is a promising tool for non-invasive modulation of cerebellar function and is under investigation for treating cerebellum-related disorders. However, its local and remote effects on sensory processing remain poorly understood. We investigated the immediate and long-term effects of Cb-tDCS on sensory-evoked responses in the cerebellum and primary somatosensory cortex (S1) of awake mice. Sensory-evoked potentials (SEPs) were recorded in Crus I/II and S1 during and after short (15 s) or long (20 min) sessions of anodal or cathodal Cb-tDCS. In addition, vGLUT1 and GAD65–67 immunoreactivity were quantified, and spectral changes in local field potentials were assessed.

Anodal and cathodal Cb-tDCS respectively induced an immediate increase and decrease in the trigeminal component in Crus I/II but no aftereffects were observed 20 min post-stimulation. In S1, Cb-tDCS resulted in polarity and intensity-dependent modulation of the N1 component during stimulation, which was opposite to the changes induced in Crus I/II, as well as a polarity-dependent modulation after stimulation. In addition, anodal Cb-tDCS was associated with reduced GAD65–67 immunoreactivity in S1, whereas vGLUT1 remained unchanged. While power spectrum analysis revealed no changes in Crus I/II, Cb-tDCS induced polarity-dependent post-stimulation changes in S1 spectral power, with higher values after cathodal stimulation.

These findings show that Cb-tDCS differentially modulates sensory processing in cerebellar and cortical circuits. While cerebellar effects are mainly transient, stimulation induces longer-lasting changes in the remote cortical area investigated, S1. This underscores the need to consider both local and distant network effects when applying Cb-tDCS in translational and clinical settings.

## Introduction

The cerebellum, traditionally associated with motor coordination and learning (Ito, 2002), has increasingly been recognized for its involvement in sensory processing and cognitive functions (Schmahmann & Sherman, 1998; Schmahmann, 2001; Ramnani, 2006; Bostan & Strick, 2018). Transcranial direct-current stimulation (tDCS) is a non-invasive brain stimulation technique that modulates neural excitability through the application of weak direct currents to the scalp (Nitsche & Paulus, 2000; Woods et al., 2016). Recent studies have demonstrated the efficacy of tDCS in modulating cortical excitability and inducing neuroplasticity in various brain regions (Stagg et al., 2018), including the cerebellum (Galea et al., 2009). Cerebellar tDCS (Cb-tDCS) has been shown to modulate motor, cognitive, and emotional behaviors by engaging distinct cerebellar circuits (Grimaldi et al., 2016). As a result, it has been suggested as a promising noninvasive neuromodulatory therapy for disorders involving cerebellar dysfunction (Manto et al., 2022). However, the specific effects of Cb-tDCS on sensory inputs remain poorly understood.

Understanding the neural mechanisms that underlie sensory processing is crucial to unraveling the complexities of perception and cognition. Previous research has implicated the cerebellum in sensory integration and processing of a wide range of sensory modalities, including visual, auditory, and somatosensory inputs (Baumann et al., 2015). However, the precise contribution of cerebellar activity to sensory processing and how it can be modulated by tDCS remains largely unexplored. Previous studies on tDCS have demonstrated its ability to modulate the amplitude of sensory-evoked potentials (SEPs) in both humans (Matsunaga et al., 2004; Dieckhöfer et al., 2006) and animals (Márquez-Ruiz et al., 2012; Sánchez-León et al., 2021). SEPs can be measured in the primary somatosensory cortex (S1) of both humans (Sugawara et al., 2015; Vaseghi et al., 2015) and rodents (Castro-Alamancos & Bezdudnaya, 2015), underscoring their importance as a valuable tool for translational studies (Sánchez-León et al., 2018). In addition to S1, SEPs can also be recorded in Crus I/II region of the cerebellum (Mapelli & D’Angelo, 2007; Roggeri et al., 2008; Márquez-Ruiz & Cheron, 2012; Fernández et al., 2021) with several components that reflect inputs from the trigeminal ganglion and cerebral cortex (Morissette & Bower, 1996; Brown & Bower, 2002). S1 is highly interconnected with the cerebellum (Schmahmann, 2001; Ramnani, 2006; Buckner et al., 2011), with information reaching the cerebellar cortex through its two inputs, from the mossy fibers through the pontine nucleus (Allen et al., 1979; Leergaard et al., 2000; Nagao, 2004; Odeh et al., 2005) and from the climbing fibers through the inferior olive (Swenson et al., 1989; Lawrenson et al., 2016). Furthermore, the cerebellum projects to S1 through the thalamus (Proville et al., 2014), closing the loop between these two areas.

Cerebellar-brain inhibition refers to a neural pathway through which the cerebellum can exert inhibitory control over other areas of the brain (Ugawa et al., 1991). This pathway involves inhibitory Purkinje cells (PCs) in the cerebellar cortex, which receive input from various sensory and motor systems. These PCs send inhibitory signals to the cerebellar nuclei (CN)—including fastigial, interposed, and dentate nuclei—the main output structures of the cerebellum (Sugihara, 2011; Kebschull et al., 2024). Subsequently, these nuclei send excitatory signals to the thalamus (Gornati et al., 2018), which then modulates the activity of different brain regions, including the cerebral cortex and basal ganglia (Bostan et al., 2013). Unraveling the mechanisms and functions of cerebellar-brain inhibition is essential for advancing our knowledge of brain function and fostering the development of effective therapies for neurological disorders.

In this study, we aim to investigate the local cerebellar effects and the downstream cortical effects in S1 of Cb-tDCS on sensory inputs in awake mice. For that, we recorded the SEPs induced by whisker stimulation in Crus I/II and S1 before, during, and after the application of short (15 s) or long (20 min) sessions of Cb-tDCS. To evaluate whether Cb-tDCS was associated with complementary neurochemical changes, we performed a post-stimulation immunohistochemical analysis of the immunoreactivity of GAD65–67 and vGLUT1 in the stimulated brain region. In addition, spectral analyses were used as complementary measures of network activity. Our findings provide valuable insights into the role of the cerebellum in sensory processing and help clarify the temporal and regional profile of Cb-tDCS effects on sensory circuits, with implications for the development of novel therapeutic strategies targeting sensory disorders.

## Material and Methods

### Animals

Experiments were conducted on adult male C57 mice (University of Seville, Spain) weighing 28–35 g. All experimental procedures were carried out in accordance with European Union guidelines (2010/63/CE) and followed the Spanish regulations (RD 53/2013 and RD 118/2021) regarding the use of laboratory animals in chronic experiments. Furthermore, these experiments were submitted to and approved by the local Ethics Committee of Pablo de Olavide University (Seville, Spain). This study was carried out in accordance with the ARRIVE guidelines.

### Surgery

Mice were prepared for the chronic recording of SEPs in Crus I/II and S1, along with simultaneous Cb-tDCS, following established protocols (Sun et al., 2020; Sánchez-León et al., 2021, 2023). The mice were initially anesthetized with a ketamine–xylazine mixture (Ketaset, 100 mg/ml, Zoetis, NJ., USA; Rompun, 20 mg/ml, Bayer, Leverkusen, Germany) at an initial dosage of 0.1 ml/20 g. Under aseptic conditions, a midline anteroposterior (AP) incision was made along the head, from the front leading edge to the lambdoid suture. The periosteum of the exposed skull surface was then removed and rinsed with saline. The animal’s head was positioned accurately to mark the bregma as the stereotaxic zero reference point. In the experiments where SEPs were recorded in Crus I/II during simultaneous Cb-tDCS, a custom-made silver ring chlorinated electrode (2.5 mm inner diameter, 3.5 mm outer diameter) was placed over the left Crus I/II region on the skull (AP = − 6 mm; Lateral = +2 mm; relative to bregma (Paxinos & Franklin, 2013)) (Fig. 1A). The electrode served as the active electrode for Cb-tDCS and was secured with dental cement (DuraLay, Ill., USA), without filling the gap between the electrode and the skull. Additionally, a hole (2 mm diameter) was drilled in the interparietal bone within the ring electrode to expose the cerebellum, and the dura mater surface was protected with wax bone (Ethicon, Johnson & Johnson, NJ., USA). In experiments where SEPs were recorded in S1 during simultaneous Cb-tDCS, the active electrode for tDCS was a polyethylene tubing (inner diameter: 2.159 mm; outer diameter: 3.251 mm; A-M Systems) placed over the stimulated region (AP = − 6 mm; Lateral = +2 mm; relative to bregma (Paxinos & Franklin, 2013)) and filled with electrogel in which the electrode from the stimulator was immersed (Fig. 5A). A hole (2 mm diameter) was drilled in the right parietal bone, centered on the right S1 vibrissa area (AP = − 0.9 mm; Lateral = − 3 mm; relative to bregma (Paxinos & Franklin, 2013)), and the dura mater surface was protected with wax bone. Furthermore, a silver electrode was implanted over the dura surface beneath the left parietal bone (AP = − 0.9 mm; Lateral = + 3 mm; relative to bregma (Paxinos & Franklin, 2013)) to serve as the electrical reference for the electrophysiological recordings. The electrode was constructed by cutting a silver wire (381 μm diameter, A-M Systems) into 1 cm pieces. A 2 mm diameter loop was formed at one end to facilitate grasping by the amplifier system, while the opposite end was braided and filed to prevent damage to the dura mater. For the histological experiments, the active electrode for Cb-tDCS consisted of a polyethylene tubing as described earlier. No trepanation was performed in the histological experiments to prevent tissue damage. Finally, a head-holding system, comprising three bolts screwed into the skull and an upside-down bolt placed over the skull perpendicular to the horizontal plane, was implanted to allow for head fixation during the experiments. The complete holding system was cemented to the skull to ensure stable head positioning.

**Figure 1.**
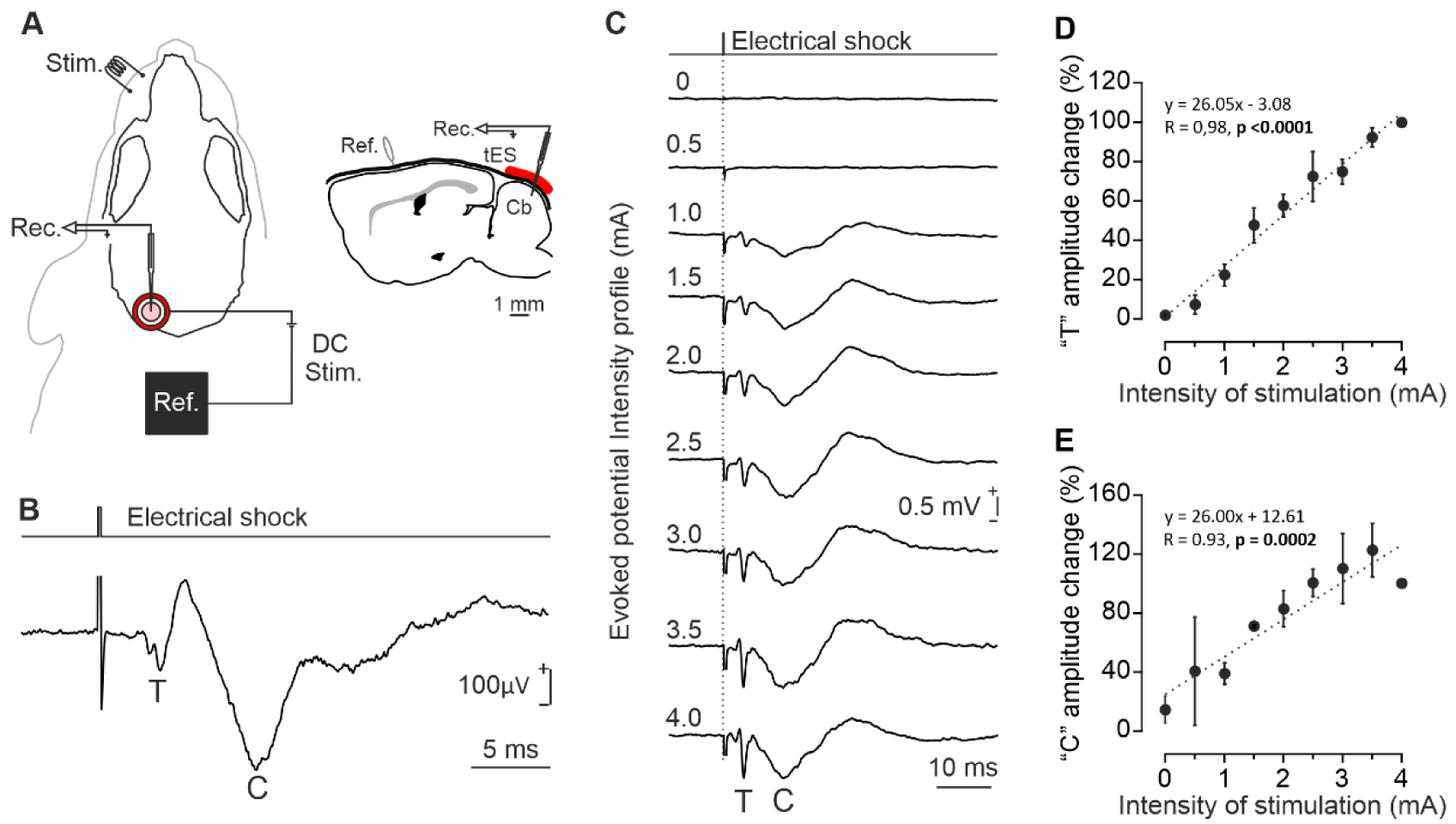
Sensory evoked potential (SEP) characterization in Crus I/II. *A*) Experimental setup for concurrent Cb-tDCS and *in vivo* electrophysiological recordings in Crus I/II. The right side shows a schematic representation of the active electrode and recording site in the lateral cerebellum. *B*) Representative profile of Crus I/II SEP induced by whisker electrical stimulation. The graph displays the different trigeminal (T) and cortical (C) components (n = 30 SEPs from a representative mouse). *C*) Intensity profile of Crus I/II-SEP. Each trace represents an average of 5 SEPs recorded at the same location but with varying intensities of whisker electrical stimulation. *D*) Quantification of amplitude changes in the T component of SEPs at different intensities of whisker electrical stimulation. Data normalized to the maximum amplitude recorded at 4 mA (N = 2 mice). E) Quantification of amplitude changes in the C component of SEPs at different intensities of whisker electrical stimulation. Data normalized to the maximum amplitude recorded at 4 mA (N = 2 mice). Cb: cerebellum; Rec.: recording; Ref.: reference; Stim.: stimulation; tES: transcranial electrical stimulation.

### Recording and stimulation procedures

Recording sessions commenced at least two days after the surgical procedure. The mice were placed on a treadmill equipped with an infrared sensor to monitor locomotion activity. The head was securely fixed to the recording table using the implanted head-holding system. To stimulate the whiskers, an electrical stimulus (0.2 ms square pulse, < 2.5 mA) was delivered through a pair of hook electrodes inserted in the left whisker pad. The electrodes were connected to an isolation unit (CS20, Cibertec, Madrid, Spain) controlled by a stimulator device (CS420, Cibertec). For characterizing the SEPs, a glass micropipette (1-5 MΩ of impedance; outer diameter: 2.0 mm; inner diameter: 1.6 mm; length: 15 cm, with inner filament; A-M Systems, WA., USA) filled with 3M NaCl was mounted on a micromanipulator (MO-10, Narishige, Tokyo, Japan) and positioned over S1 area. The electrical stimulus was delivered to the whisker pad every 10 ± 2 s while the micropipette was gradually lowered and the current intensity adjusted (0.7 - 2.5 mA) until the maximum amplitude SEP was achieved. Subsequently, the current intensity of the whisker electrical pulses was reduced to elicit a SEP with half of the maximum amplitude. This allowed for the observation of changes in the components of the SEP during and after Cb-tDCS intervention. All recordings were acquired using an amplifier (BVC-700A, Dagan corporation, MN., USA) connected to a dual extracellular-intracellular headstage (8024, Dagan corporation; gain error ± 1%, noise 4.5 µV root mean square). The sampling rate was set at 25 kHz, and the amplitude resolution was 12 bits (CED Micro 1401; Cambridge Electronic Design, Cambridge, UK).

### Transcranial electrical stimulation

The different protocols for transcranial currents were designed in Spike2 software (Cambridge Electronic Design (CED), Cambridge, U.K.) and transmitted to a battery-powered linear stimulus isolator (WPI A395, Fl., USA). Cb-tDCS was applied between the ring electrode (in the experiments combining SEP recording in Crus I/II) or the plastic tubing (in experiments combining SEP recording in S1) placed over Crus I/II. A reference electrode consisting of a rubber rectangle (6 cm^2^) moistened with electrogel (Electro-Cap International, OH., USA) was attached to the back of the mouse. A silver wire was inserted into the rubber electrode for connection to the stimulator. To assess the immediate effects induced by Cb-tDCS, 15 s pulses of anodal and cathodal tDCS were applied with a 10-second gap of non-stimulation between them. Cb-tDCS was delivered at 200 μA to assess local effects in the cerebellum and at 200 and 300 μA to evaluate changes in S1. For assessing after-effects changes in Crus I/II, Cb-tDCS was delivered for 20 min at 200 μA for cathodal stimulation, 20 min at 200 μA for anodal stimulation, and for 30 s at 200 μA anodal for sham stimulation. To evaluate after-effects changes in S1, Cb-tDCS was administered for 20 min at 300 μA for cathodal stimulation, 20 min at 300 μA for anodal stimulation, and for 30 s at 300 μA anodal for sham stimulation. This intensity was selected because, in the initial experiments assessing immediate effects in S1, 300 μA produced larger and more robust modulation of SEP amplitude than 200 μA, whose effects were weaker and less consistent. The current was directly monitored during experiments to ensure that it matched the desired level indicated in the stimulator.

### Histology

To examine potential histological changes in Crus I/II and S1 associated with Cb-tDCS, a separate group of animals received 20 min of anodal, cathodal, or sham Cb-tDCS at 300 μA. Fifteen min after cessation of Cb-tDCS, mice were deeply anesthetized with a ketamine–xylazine mixture (Ketaset, 100 mg/ml; Rompun, 20 mg/ml) and transcardially perfused with 0.9% saline, followed by 4% paraformaldehyde (PanReac, Barcelona, Spain) in PBS. The brains were then extracted and stored in 4% paraformaldehyde for 24 hours. Subsequently, they were cryoprotected in 30% sucrose in PB for the next 48 hours and then sectioned into 50 μm coronal slices using a freezing microtome (CM1520, Leica, Wetzlar, Germany). After three 10-min washes with PBS, sections were blocked with 10% Normal Donkey Serum (NDS, 566460, Merck, Darmstadt, Germany) in PBS containing 0.2% Triton X-100 (Sigma-Aldrich, Mo., USA) (PBS-Tx-10% NDS). They were then incubated overnight at room temperature in darkness with mouse anti-vesicular Glutamate Transporter 1 (vGLUT1, 1:1000, MAB5502, Merck) or rabbit anti-Glutamate Decarboxylase 65-67 (GAD65-67, 1:1000, AB1511, Merck). After three washes, the sections were incubated for 1 hour at room temperature in darkness with appropriate secondary antibodies: Alexa Fluor 488 donkey anti-mouse IgG (H+L) (1:400, A21202, Thermo Fisher Scientific, Mass., USA) or Alexa Fluor 555 donkey anti-rabbit IgG (H+L) (1:400, A31572, Thermo Fisher Scientific) in PBS-Tx-5% NDS. Following three washes with PBS, the sections were mounted on glass slides and coverslipped. Confocal images were acquired using a Nikon A1R HD25 confocal microscope (Nikon, Tokyo, Japan) in sequential scanning mode with a 63× oil-immersion objective (NA 1.30). Alexa Fluor 488 and Alexa Fluor 555 were excited using 488 and 561 nm laser lines, respectively, and the corresponding emission signals were collected using 500–550 nm and 570–620 nm detection windows. Z-series of optical sections were acquired at 0.5 μm intervals, and images collected at a resolution of 1024 × 1024 pixels (0.17067 μm/pixel). Images used for analysis were obtained at depths ranging from 30.5 to 45.5 μm. Acquisition settings were kept constant across groups.

### Data analysis

SEP amplitude was computed using the peak-to-peak command in Spike2 software, where the maximum negative voltage value (N1) was subtracted from the maximum positive voltage value (P1) of the preceding peak. SEPs recorded when the animal was running were excluded from the analysis, as well as potentials containing electrical artifacts.

Confocal images were processed using Fiji software (http://fiji.sc/Fiji) with a custom-built macro. The fluorescence background was subtracted, and five square regions of interest (ROIs) measuring 100 × 100 pixels (291.31 μm²) were selected in comparable areas of the same anatomical region across animals, avoiding nuclei and nonspecific structures such as blood vessels. Particle analysis was not performed at the level of individual cells, but within these standardized ROIs. Each image within the ROI was converted to binary, and the “Analyze Particles” command was used to count and measure aggregates of vGLUT1 and GAD65-67. The same acquisition and Fiji/ImageJ analysis criteria were applied across samples, and the particle counts obtained from the selected ROIs were averaged to obtain one representative value per hemisphere per animal.

To analyze the spectral dynamics of the neural oscillations using the Fast Fourier Transform (FFT) an analysis of the *induced activity* was performed. The average SEP from each subject was subtracted from each condition, temporal period, and subject (Bastiaansen & Hagoort, 2003; Tallon-Baudry & Bertrand, 1999). Power spectrum computation was carried out using the FieldTrip toolbox (Oostenveld et al., 2011), employing the fast Fourier transform (multitaper method) for frequencies ranging from 3 to 300 Hz. Power spectrum values were extracted from each trial and averaged for each animal and temporal period. The calculated power was referenced to power obtained during the control period, resulting in the ratio of the power spectrum during each temporal period relative to the control period for each subject. Spectral power was then determined for specific power bands: theta (4-10 Hz), beta (10-30 Hz), low gamma (30-45 Hz), and high gamma (60-100 Hz). These ratios were averaged within each group. Power bands from 20 min post-tDCS were compared to the control period.

Statistical analysis was conducted using SigmaPlot 11.0 (Systat Software Inc, San Jose, CA., USA), IBM SPSS version 31 (IBM, Armonk, NY) and Matlab 2015a (MathWorks Inc.). Normality was assessed using the Shapiro–Wilk test (P value > 0.05). For immediate-effect experiments, amplitude values recorded during tDCS were normalized to the corresponding control period and expressed as percentage of control (control = 100%), whereas latency values were expressed as differences relative to control (control = 0 ms). SEP data were analyzed using linear mixed-effects models with animal as a random intercept. For immediate effects, the fixed factors were COMPONENT, TIME, and POLARITY in Crus I/II, and INTENSITY, TIME, and POLARITY in S1. For after-effect analyses in both Crus I/II and S1, the fixed factors were TIME and POLARITY. When appropriate, post hoc pairwise comparisons were performed using Bonferroni correction. Immunohistochemical data were analyzed separately for each marker and region using linear mixed-effects models with POLARITY (anodal, cathodal, sham) and HEMISPHERE (stimulated vs non-stimulated) as fixed factors, and animal as a random intercept. Spectral power data from Crus I/II and S1 were analyzed separately using linear mixed-effects models with POLARITY (anodal vs cathodal), PERIOD (control vs post-tDCS), and BAND as fixed factors, and animal as a random intercept. When appropriate, post hoc pairwise comparisons were performed using Bonferroni correction. The results are presented as mean ± SEM, and statistical significance was set at p < 0.05 for all analyses.

## Results

### Characterization of cerebellar SEPs in response to whisker stimulation

To assess potential changes in neuronal excitability within the Crus I/II region of the cerebellum during and after tDCS, SEPs in response to whisker stimulation were chronically recorded in awake, head-restrained mice (N = 14; Fig. 1A). Electrical stimulation of the whisker evoked an ipsilateral short-latency SEP in the Crus I/II, characterized by two prominent negative waves corresponding to trigeminal (T) and cortical (C) responses, peaking at 3.79 ± 0.69 ms and 12.57 ± 1.12 ms, respectively (Fig. 1B). The amplitudes of these SEP components demonstrated a linear relationship with the intensity of the electrical stimuli applied to the whiskers, as depicted in Figure 1C. The correlation coefficients for the T and C components were R = 0.98 (p < 0.0001, N = 2) and R = 0.93 (p = 0.0002, N = 2), respectively (Fig. 1D,E). During the experiments, the final current intensity applied to the whiskers was adjusted to elicit a T component with half of the maximum amplitude, allowing for modulation of the SEP amplitude during and after Cb-tDCS intervention.

### Cb-tDCS modulates the amplitude of SEPs recorded in the cerebellum during its application but does not cause observable long-term effects

To test the immediate effects of Cb-tDCS on Crus I/II excitability we recorded SEPs induced by whisker pad stimulation during simultaneous short-duration (15 s, including 5 s ramp up and 5 s ramp down) anodal and cathodal tDCS pulses (± 200 µA) (Fig. 2A). The calculated density current for the used tDCS intensity was 4.26 mA/cm^2^. The peak-to-peak amplitudes and the latencies of the T and C components of SEPs recorded just before Cb-tDCS pulses were compared with those recorded during Cb-tDCS application. Figure 2A shows the averaged SEPs (n = 15) during anodal (dark red trace) and cathodal (dark blue trace) Cb-tDCS, as well as the control conditions for anodal (light red trace) and cathodal (light blue trace), for a representative animal. Mean data obtained from the group of animals participating in the experiment (N = 10) for T and C components are represented in Figure 2B,C. Data were normalized to the values during the control condition for anodal or cathodal stimulation. To assess the immediate effects of Cb-tDCS on cerebellar SEPs, amplitude data were analyzed using a linear mixed-effects model including COMPONENT (T vs C), TIME (control vs during stimulation), and POLARITY (anodal vs cathodal) as fixed factors, and animal as a random intercept. This analysis revealed a significant COMPONENT × POLARITY interaction (F_1,63_ = 12.993, p < 0.001) and a significant COMPONENT × TIME × POLARITY interaction (F_1,63_ = 12.993, p < 0.001), whereas no significant main effects of COMPONENT, TIME, or POLARITY were found (all p > 0.3). Descriptively, anodal Cb-tDCS increased the amplitude of the T component to 111.2 ± 3.61% of control, whereas cathodal Cb-tDCS decreased it to 86.13 ± 3.96%. For the C component, anodal stimulation reduced amplitude to 88.57 ± 7.38% of control, whereas cathodal stimulation produced values of 103.2 ± 6.21%. Post hoc pairwise comparisons with Bonferroni correction showed that anodal Cb-tDCS significantly increased the amplitude of the T component relative to control (p = 0.047), while significantly decreasing the amplitude of the C component (p = 0.042). Cathodal Cb-tDCS significantly decreased the amplitude of the T component relative to control (p = 0.014), whereas the C component did not show significant changes (p = 0.562). Within each stimulation polarity, the T and C components also differed significantly from each other during stimulation, both for anodal (p < 0.001) and cathodal Cb-tDCS (p = 0.003). In addition, during stimulation, anodal and cathodal Cb-tDCS differed significantly for both the T component (p < 0.001) and the C component (p = 0.010). These results indicate that the immediate effects of Cb-tDCS in Crus I/II are component- and polarity-dependent. Latency data were also analyzed using a linear mixed-effects model with COMPONENT (T vs C), TIME (control vs during stimulation), and POLARITY (anodal vs cathodal) as fixed factors, and animal as a random intercept. No significant main effects of COMPONENT, TIME, or POLARITY, and no significant interactions among factors were detected (all p > 0.28), indicating that immediate Cb-tDCS did not induce reliable changes in the latency of SEP components recorded in Crus I/II. In summary, Cb-tDCS induced polarity-dependent changes in the amplitude of cerebellar SEPs during stimulation, with the most robust modulation observed in the trigeminal component, whereas no significant effects were found on SEP latency.

**Figure 2.**
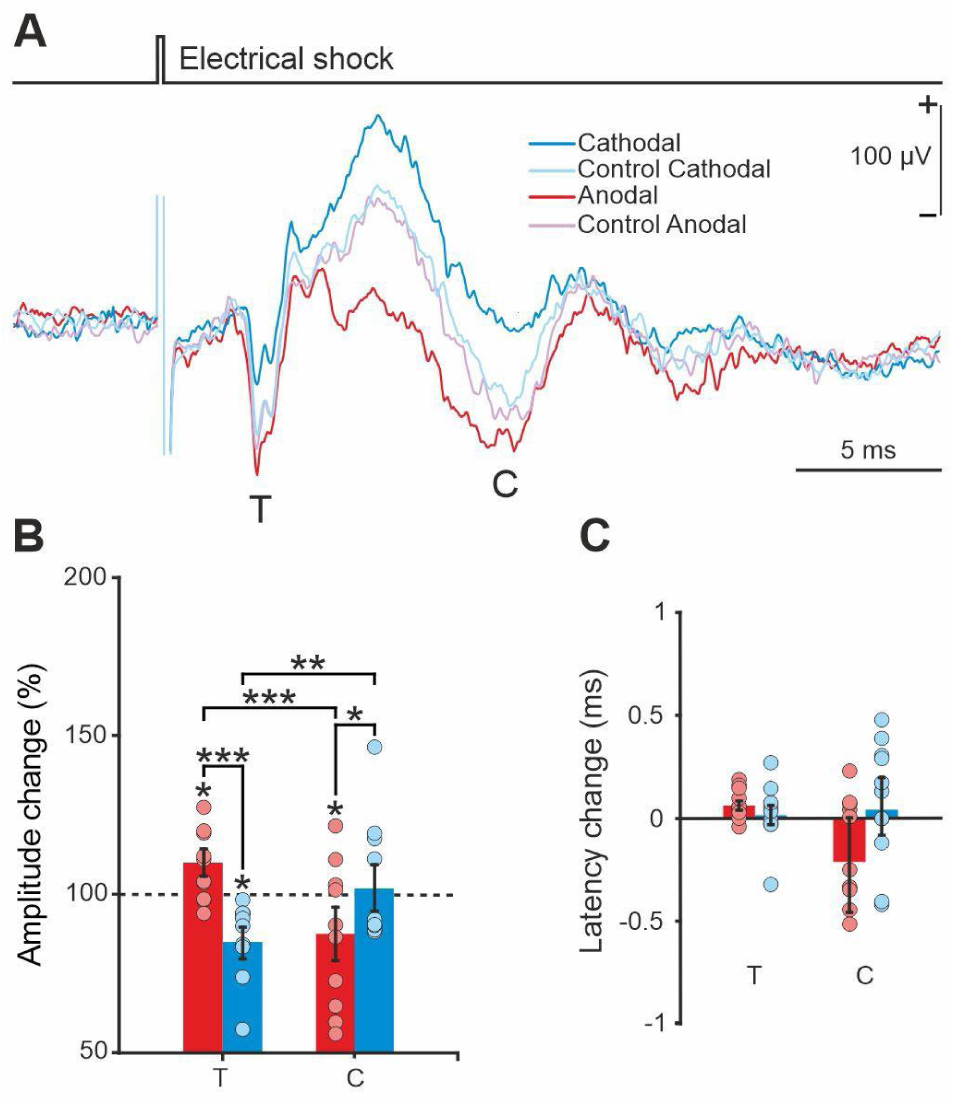
Immediate effects of Cb-tDCS over SEPs in Crus I/II region. *A*) Average SEP (n = 15) recorded in Crus I/II from a representative animal during control before anodal stimulation (light-red trace), anodal (red trace), control before cathodal (light-blue trace), and cathodal (blue trace) Cb-tDCS applied at 4.26 mA/cm^2^. *B*) Quantification and statistical analysis of Cb-tDCS effects on the amplitude of the trigeminal (T) and cortical (C) components of Crus I/II SEP. The mean (bars) and individual amplitude data (circles) are represented as percentage of change compared to control values for all animals (N = 10 mice). Statistical analysis was performed using a linear mixed-effects model with COMPONENT, TIME, and POLARITY as fixed factors and animal as a random intercept, followed by Bonferroni-corrected pairwise comparisons, with *p < 0.05, **p < 0.01, ***p < 0.001. *C*) Quantification and statistical analysis of the effects of Cb-tDCS on the latency of SEP-components. The mean difference from control values for all animals is represented. No significant effects on latency were detected. Error bars indicate the standard error of the mean (SEM).

**Figure 3.**
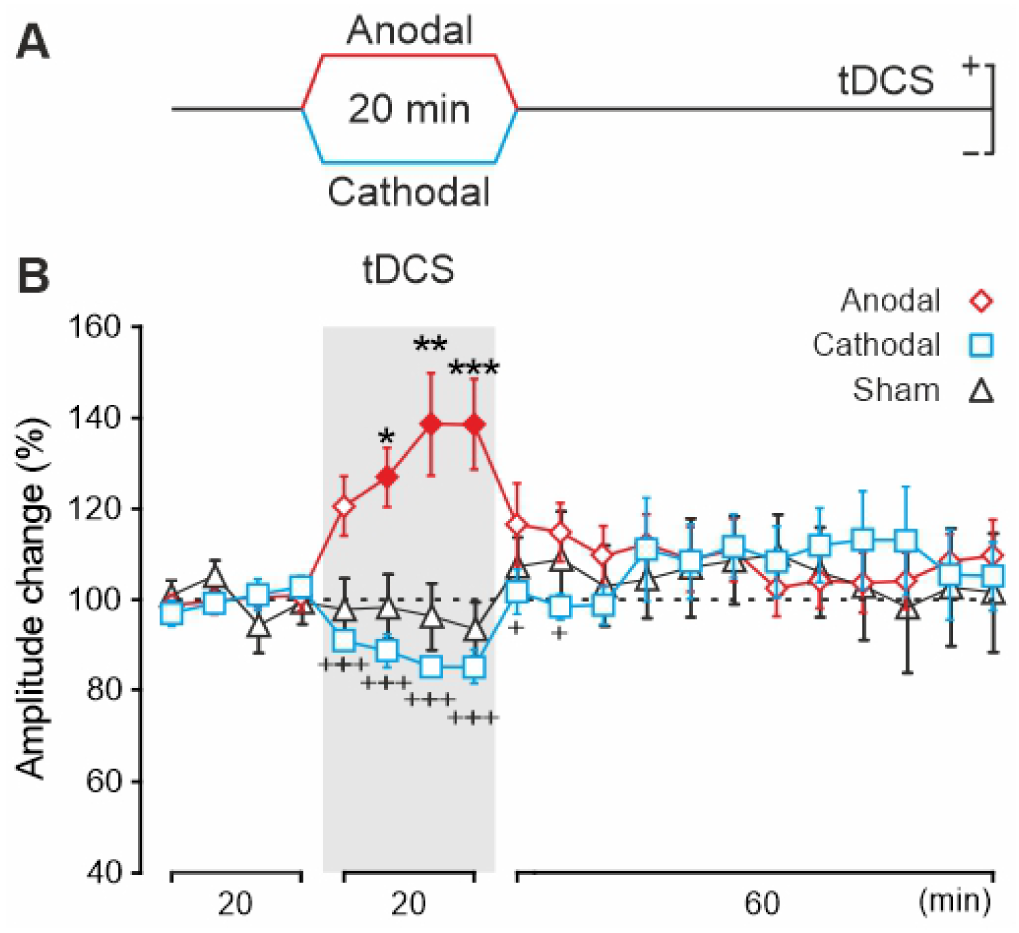
Cb-tDCS after-effects over SEPs in Crus I/II region. *A*) Schematic representationof the Cb-tDCS protocol applied for long-term experiments (20 min, 4.26 mA/cm^2^). *B*) Normalized amplitude change of the trigeminal (T) component, averaged every 5 min for 20 min of anodal (red diamonds), cathodal (blue squares), or sham (black triangles) tDCS. Statistical analysis was performed using a linear mixed-effects model with POLARITY and TIME as fixed factors and animal as a random intercept, followed by Bonferroni-corrected pairwise comparisons. Filled red diamonds indicate significant differences compared to the last control period (N = 12 mice for anodal, N = 11 for cathodal, N = 7 mice for sham, p < 0.05). Asterisks indicate statistical differences between anodal and sham at the corresponding time bin (N = 12 mice for anodal, N = 11 for cathodal, N = 7 mice for sham, *p < 0.05; **p < 0.01; ***p < 0.001). Crosses indicate statistical differences between anodal and cathodal at the corresponding time bin (^+^p < 0.05; ^+++^p < 0.001). Error bars indicate the standard error of the mean (SEM).

To test the potential after-effects of Cb-tDCS over Crus I/II excitability we recorded SEPs induced by whisker pad stimulation (every 10 ± 2 s) in three different randomly assigned experimental conditions, anodal (N = 12), cathodal (N = 11) or sham (N = 7) group. During experimental sessions, SEPs were recorded for 20 min before Cb-tDCS, during continuous anodal (200 µA, 20 min), cathodal (-200 µA, 20 min) or sham (200 µA, 30 s) Cb-tDCS, and for 1 hour after Cb-tDCS. Every 5-minute interval SEP waveforms were averaged and normalized with respect to baseline values (control condition, before Cb-tDCS). To assess the after-effects of Cb-tDCS on the trigeminal SEP component recorded in Crus I/II, amplitude values were analyzed using a linear mixed-effects model with POLARITY (anodal, cathodal, sham) and TIME (each 5-minutes averages) as fixed factors, and animal as a random intercept. The analysis revealed a significant main effect of POLARITY (F_2,34.486_ = 33.065, p < 0.001) and a significant POLARITY × TIME interaction (F_38,520.538_ = 3.649, p < 0.001), whereas the main effect of TIME was not significant (F_19,520.538_ = 0.927, p = 0.549). Descriptively, anodal Cb-tDCS increased the normalized amplitude of the trigeminal component, reaching values of up to 138.58 ± 9.86% (N = 12) of control during stimulation, whereas cathodal Cb-tDCS reduced it to values as low as 85.4 ± 3.68% (N = 11). Bonferroni-corrected pairwise comparisons showed that, relative to its own baseline, anodal Cb-tDCS produced a significant increase in trigeminal SEP amplitude during the 5–10 min (p = 0.028), 10–15 min (p < 0.001), and 15–20 min (p < 0.001) intervals after stimulation onset. In addition, during the 20 min stimulation period, anodal Cb-tDCS produced significantly higher amplitudes than cathodal stimulation at all 5-min bins (all p < 0.001), and significantly higher amplitudes than sham during the 5–10, 10–15, and 15–20 min intervals after stimulation onset. In contrast, after stimulation cessation, no consistent differences relative to sham were detected, and differences between anodal and cathodal conditions were restricted to the first 10 min of the post-stimulation period (all p < 0.05). These results indicate that the effects of Cb-tDCS on the trigeminal component in Crus I/II were mainly restricted to the stimulation period and did not persist robustly after stimulation offset. No consistent changes were detected in the cortical component across stimulation conditions. Latency analysis did not reveal reliable long-term effects of Cb-tDCS on SEP timing in Crus I/II.

Finally, to explore potential molecular changes induced by Cb-tDCS, we used antibodies against vGLUT1 and GAD65-67 to assess possible modifications of the excitation/inhibition balance in the transcranially stimulated Crus I/II region. A group of animals prepared for Cb-tDCS application during whisker stimulation (no electrophysiological recordings were carried out in this experiment) was randomly assigned to anodal (N = 7), cathodal (N = 7) or sham (N = 7) conditions. Representative confocal images from the non-stimulated right hemisphere and the transcranially stimulated left hemisphere are presented in Figure 4 for sham (at top), anodal (at middle), and cathodal (at bottom) groups for GAD65-67 (Fig. 4A) and vGLUT1 (Fig. 4B). The number of GAD65-67 and vGLUT1 positive clusters of puncta in the stimulated and non-stimulated Crus I/II region were analyzed in the cathodal, anodal and sham groups.

**Figure 4.**
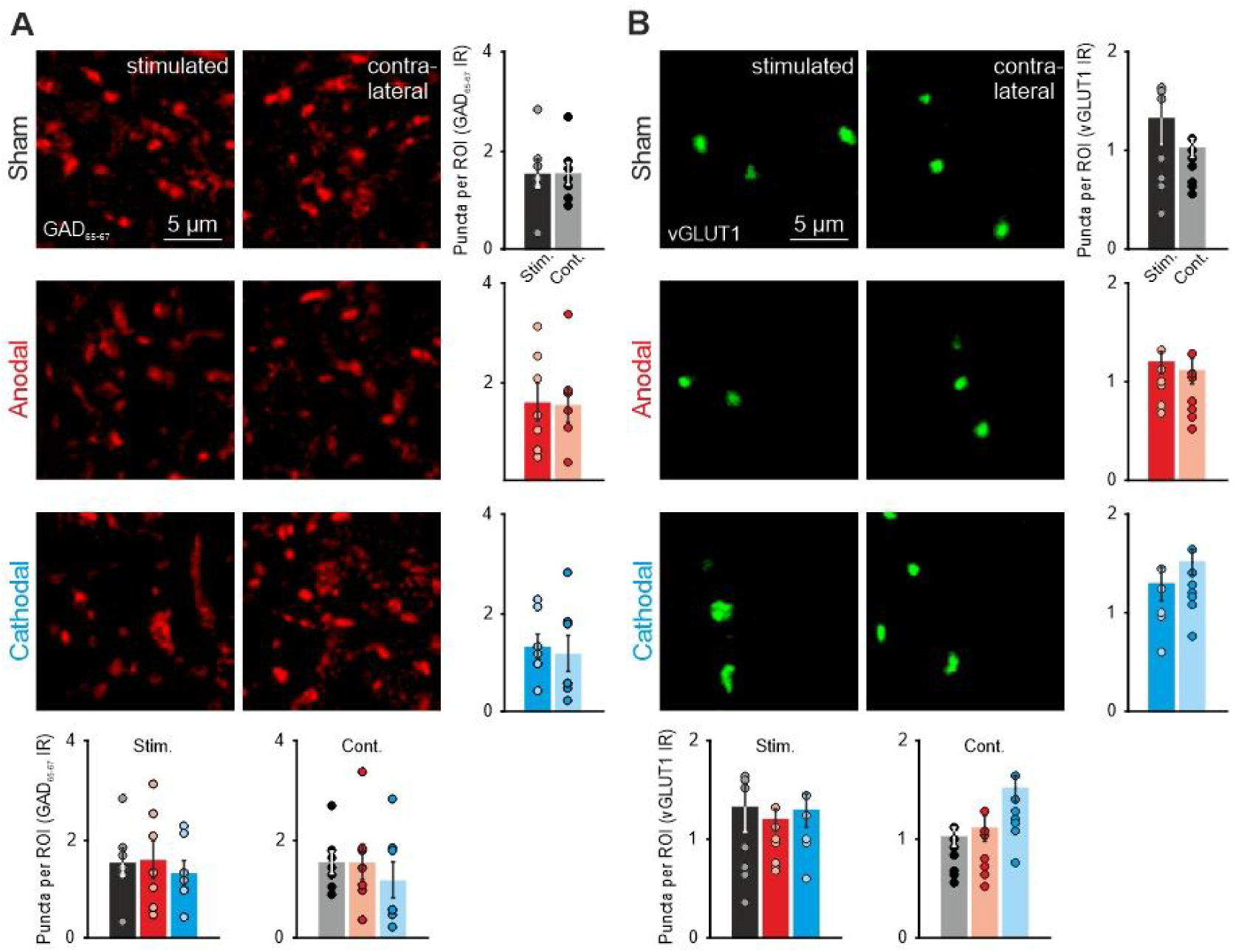
Immunohistochemical immunoreactivity in Crus I/II after 20 min of Cb-tDCS. *A,B*) Confocal photomicrographs, quantification, and statistical analysis of GAD65-67 (*A*) or vGLUT1 immunoreactivity (*B*) in Crus I/II after 20 min of sham condition (upper row, N = 7 mice), anodal Cb-tDCS (middle row, N = 7 mice) and cathodal Cb-tDCS (lower row, N = 7 mice). Bars represent mean ± SEM and circles represent individual animals. Bottom graphs show the quantification for each hemisphere across stimulation conditions. Statistical analysis was performed using linear mixed-effects models with POLARITY and HEMISPHERE as fixed factors and animal as a random intercept. No significant effects were detected for either marker. GAD65-67: Glutamic acid decarboxylase isoforms 65 and 67; vGLUT1: vesicular glutamate transporter 1; ROI: region of interest.

**Figure 5.**
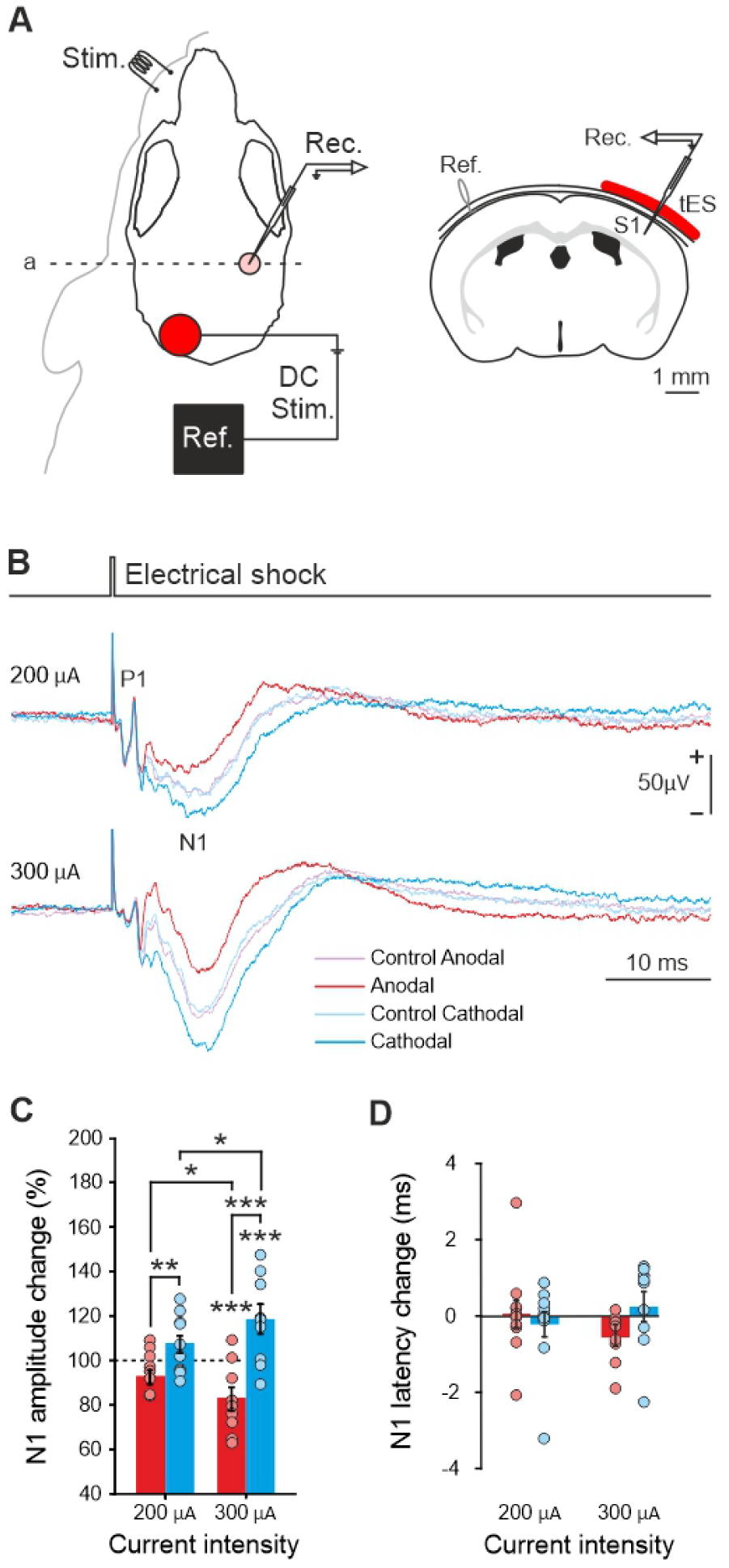
Immediate effects of Cb-tDCS over SEPs in S1. *A*) Experimental setup for concurrent Cb-tDCS and *in vivo* electrophysiological recordings in S1. A schematic coronal representation of the active electrode and recording site in the somatosensory cortex is shown on the right. *B*) Average SEP (n = 15) recorded in S1 from a representative animal during control before anodal stimulation (light-red trace), anodal (red trace), control before cathodal (light-blue trace), and cathodal (blue trace) Cb-tDCS applied at 5.46 mA/cm^2^ (upper trace) and 8.19 mA/cm^2^ (lower trace). *C*) Quantification and statistical analysis of Cb-tDCS effects on the amplitude of the N1 component of S1 SEP. The mean (bars) and individual amplitude data (circles) are represented as percentage of change compared to control values for all animals (N = 11 mice for 5.46 mA/cm^2^, N = 9 for 8.19 mA/cm^2^). Statistical analysis was performed using a linear mixed-effects model with INTENSITY, TIME, and POLARITY as fixed factors and animal as a random intercept, followed by Bonferroni-corrected pairwise comparisons, with *p < 0.05, **p < 0.01, ***p < 0.001. *D*) Quantification of latency changes in the N1 component of the S1 SEP during anodal and cathodal Cb-tDCS at 200 and 300 µA. No significant effects on latency were detected. The mean difference from control values for all animals is represented. Error bars indicate the standard error of the mean (SEM).

**Figure 6.**
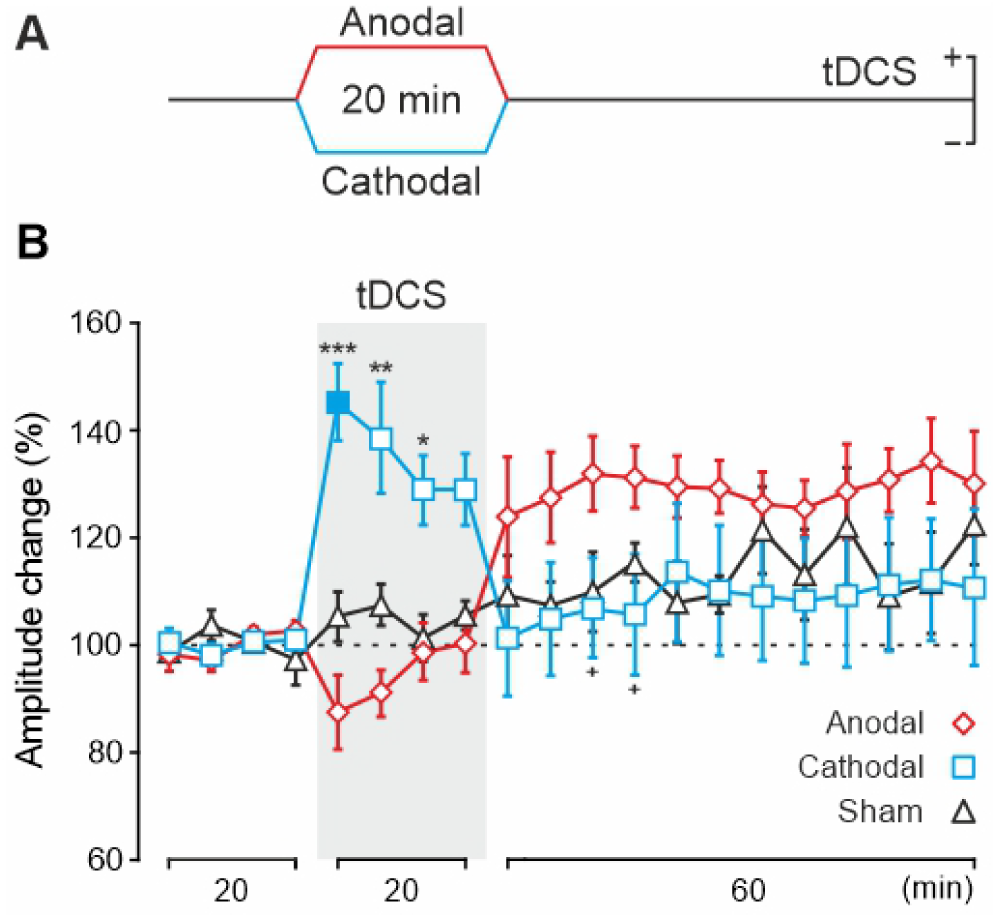
Cb-tDCS after-effects over SEPs in S1. *A*) Illustration of the Cb-tDCS protocol applied for long-term experiments (20 min, 4.26 mA/cm^2^). *B*) Normalized amplitude changes of the N1 component of the S1 SEP, averaged every 5 min for 20 min of anodal (red diamonds), cathodal (blue squares), or sham (black triangles) Cb-tDCS. Statistical analysis was performed using a linear mixed-effects model with POLARITY and TIME as fixed factors and animal as a random intercept, followed by Bonferroni-corrected pairwise comparisons. Filled blue square indicate statistical differences compared to the last control period (p < 0.05). Asterisks indicate statistical differences between the same temporal period for anodal or cathodal Cb-tDCS compared to sham (N = 5 mice for anodal, N = 5 for cathodal, N = 8 mice for sham, *p < 0.05; **p < 0.01; ***p < 0.001, Bonferroni). Crosses indicate statistical differences between anodal and cathodal at the corresponding time bin (+p < 0.05). Error bars indicate the standard error of the mean (SEM).

To examine whether a single session of Cb-tDCS induced persistent histological changes in Crus I/II, vGLUT1 and GAD65-67 immunoreactivity were analyzed using linear mixed-effects models with POLARITY (anodal, cathodal, sham) and HEMISPHERE (stimulated vs non-stimulated) as fixed factors, and animal as a random intercept. For vGLUT1, no significant main effects of POLARITY (F_2,18_ = 1.549, p = 0.239) or HEMISPHERE (F_1,18_ = 0.223, p = 0.642), and no significant POLARITY × HEMISPHERE interaction (F_2,18_ = 1.816, p = 0.191) were detected. Likewise, for GAD65-67, no significant main effects of POLARITY (F_2,18_ = 0.619, p = 0.549) or HEMISPHERE (F_1,18_ = 0.075, p = 0.787), and no significant POLARITY × HEMISPHERE interaction (F_2,18_ = 0.027, p = 0.973) were observed. These results indicate that a single session of Cb-tDCS did not induce detectable persistent changes in glutamatergic or GABAergic immunoreactivity in Crus I/II. This result is consistent with the absence of long-term effects described in the previously mentioned electrophysiological experiments.

### Cb-tDCS induces immediate and after-effects on SEPs recorded in S1

To assess the potential impact of Cb-tDCS on cerebellar-brain inhibition processes, we recorded the SEPs in S1 elicited by whisker pad stimulation (Fig. 5A). Recordings were conducted during simultaneous short-duration (15 s) and throughout the 20-minute period of both anodal and cathodal Cb-tDCS pulses, as well as during the post-stimulation period. Electrical whisker stimulation evoked a contralateral short-latency SEP in the vibrissa S1 area (Fig. 5B) consisting of a first positive component (P1) peaking at 3.18 ± 0.10 ms (N = 8), followed by a negative wave (N1) at 11.73 ± 1.72 ms (N = 8).

To test the immediate effects of Cb-tDCS on S1 excitability we recorded SEPs induced by whisker pad stimulation during simultaneous short-duration (15 s, including 5 s ramp up and 5 s ramp down) anodal and cathodal Cb-tDCS pulses at two different intensities (± 200 and ± 300 µA, equivalent to 5.46 and 8.19 mA/cm^2^, respectively). SEPs recorded in S1 just before Cb-tDCS pulses were used as controls to normalize the peak-to-peak amplitude of N1 component in both anodal and cathodal conditions. Figure 5B shows the averaged SEPs (n = 30) during anodal (dark red trace), cathodal (dark blue trace) Cb-tDCS, and the control conditions for anodal (light red trace) and cathodal (light blue trace) at ± 200 and ± 300 µA from a representative animal. Mean data obtained from the groups of animals participating in the experiment (200 µA, N = 11; 300 µA, N = 9) are represented in figure 5C,D (data was normalized by the values during control condition for anodal or cathodal stimulation). To assess the immediate effects of Cb-tDCS on S1 excitability, SEP amplitudes were analyzed using a linear mixed-effects model with INTENSITY (200 vs 300 µA), TIME (control vs during stimulation), and POLARITY (anodal vs cathodal) as fixed factors, and animal as a random intercept. The analysis revealed a significant main effect of POLARITY (F_1,72_ = 28.919, p < 0.001), as well as significant INTENSITY × POLARITY (F_1,72_ = 4.925, p = 0.030), TIME × POLARITY (F_1,72_ = 28.919, p < 0.001), and INTENSITY × TIME × POLARITY (F_1,72_ = 4.925, p = 0.030) interactions. No significant main effects of INTENSITY or TIME alone were detected (both p > 0.89). Descriptively, at 200 µA, anodal Cb-tDCS reduced SEP amplitude to 92.51 ± 3.36% of control, whereas cathodal stimulation increased it to 107.41 ± 3.75%. At 300 µA, the effect was stronger, with anodal stimulation reducing SEP amplitude to 82.69 ± 5.27% of control and cathodal stimulation increasing it to 118.3 ± 6.61 %. Bonferroni-corrected pairwise comparisons showed that at 300 µA, anodal stimulation significantly reduced SEP amplitude relative to control (p < 0.001), whereas cathodal stimulation significantly increased it (p < 0.001). At 200 µA, the changes did not reach statistical significance (anodal: p = 0.099; cathodal: p = 0.102). In addition, during stimulation, SEP amplitudes differed significantly between 200 and 300 µA for both anodal (p = 0.041) and cathodal (p = 0.021) stimulation, and anodal and cathodal differed significantly at both 200 µA (p=0.001) and 300 µA (p < 0.001). These results indicate that the immediate effects of Cb-tDCS on S1 are polarity-dependent and become more pronounced at higher stimulation intensity. Latency data were also analyzed using a linear mixed-effects model with INTENSITY, TIME, and POLARITY as fixed factors, and animal as a random intercept. No significant main effects or interactions were detected (all p > 0.11), indicating that immediate Cb-tDCS did not induce reliable changes in SEP latency in S1. In summary, immediate Cb-tDCS induced polarity-dependent modulation of S1 SEP amplitude in the opposite direction to that observed in Crus I/II, and intensity-dependent modulation with stronger effects at 300 µA, whereas no significant effects were found on SEP latency. Since short-term effects observed for 200 µA stimulation were weaker and less reliable than during 300 µA, we decided to apply 300 µA for the subsequent experiments.

To test the potential after-effects of Cb-tDCS on S1 excitability, we recorded SEPs induced by whisker pad stimulation (every 10 ± 2 s) in three different randomly assigned experimental conditions, anodal (N = 5), cathodal (N = 5) or sham (N = 8) groups. During experimental sessions, SEPs were recorded in S1 for 20 min before Cb-tDCS, during continuous anodal (+ 300 µA, 20 min), cathodal (- 300 µA, 20 min) or sham (300 µA, 30 s) Cb-tDCS, and for 1 hour after Cb-tDCS. Every 5-minute interval SEP waveforms were averaged and normalized with respect to baseline values (control condition, before Cb-tDCS). Amplitude values were analyzed using a linear mixed-effects model with POLARITY (anodal, cathodal, sham) and TIME (each 5-minutes averages) as fixed factors, and animal as a random intercept. The analysis revealed a significant main effect of TIME (F_19,289.118_ = 3.179, p < 0.001) and a significant POLARITY × TIME interaction (F_38,289.118_ = 3.251, p < 0.001), whereas the main effect of POLARITY was not significant (F_2,18.807_ = 2.143, p = 0.145). Descriptively, during stimulation, anodal Cb-tDCS was associated with lower SEP amplitudes, reaching 87.45 ± 7.21% of control during the first 5 min after stimulation onset, whereas cathodal stimulation was associated with higher amplitudes, reaching 145.22 ± 6.96% during the same interval. Bonferroni-corrected pairwise comparisons showed that, relative to its own baseline, anodal Cb-tDCS produced a significant increase in SEP amplitude during the 0–5 min (p = 0.004) interval after stimulation onset. In addition, cathodal Cb-tDCS induced significantly higher amplitudes than anodal stimulation across all 5-min bins of the 20-min stimulation period (all p ≤ 0.019), and significantly higher amplitudes than sham during the first 15 min of stimulation (all p ≤ 0.026). During the post-stimulation period, differences between groups were more limited, with anodal amplitudes exceeding cathodal values 10–15 min (p = 0.049) and 15–20 min (p = 0.043) after stimulation offset, whereas no consistent differences relative to sham were detected. Latency values were also analyzed, and no significant effects of Cb-tDCS were detected. These results indicate that Cb-tDCS produced polarity-dependent temporal dynamics in S1, with the clearest differences occurring during stimulation and only limited divergence between polarities after stimulation offset, whereas no significant effects were detected on SEP latency.

Finally, we used antibodies against vGLUT1 and GAD65-67 to assess possible modifications of the excitation/inhibition balance in S1 after transcranial stimulation of the distant Crus I/II region. A group of animals prepared for Cb-tDCS application during whisker stimulation (no electrophysiological recordings were carried out in this experiment) was randomly assigned to anodal (N = 7), cathodal (N = 7), or sham (N = 7) conditions. Representative confocal images from the left and right S1 hemispheres, including the hemisphere receiving cerebellar transcranial stimulation inputs, are presented in Figure 7 for the sham group (at the top), anodal group (in the middle), and cathodal group (at the bottom) for GAD65-67 (Fig. 7A) and vGLUT1 (Fig. 7B). The number of GAD65-67 and vGLUT1 positive clusters of puncta in both S1 regions was analyzed as previously. For vGLUT1, the linear mixed-effects model revealed no significant main effects of POLARITY (F_2,18_ = 0.973, p = 0.397) or HEMISPHERE (F_1,18_ = 0.962, p = 0.340), and no significant POLARITY × HEMISPHERE interaction (F_2,18_ = 0.025, p = 0.975). In contrast, for GAD65-67, a significant main effect of POLARITY was observed (F_2,18_ = 3.777, p = 0.043), whereas neither HEMISPHERE (F_1,18_ = 0.007, p = 0.935) nor the POLARITY × HEMISPHERE interaction (F_2,18_ = 2.010, p = 0.163) reached significance. Bonferroni-corrected pairwise comparisons showed that GAD65-67 immunoreactivity was significantly lower in the anodal group than in the sham group (p = 0.040), whereas the remaining comparisons were not significant (anodal vs cathodal, p = 0.511; cathodal vs sham, p = 0.610). These findings indicate that anodal Cb-tDCS was associated with reduced GAD65-67 immunoreactivity in S1, while vGLUT1 immunoreactivity remained unchanged.

**Figure 7.**
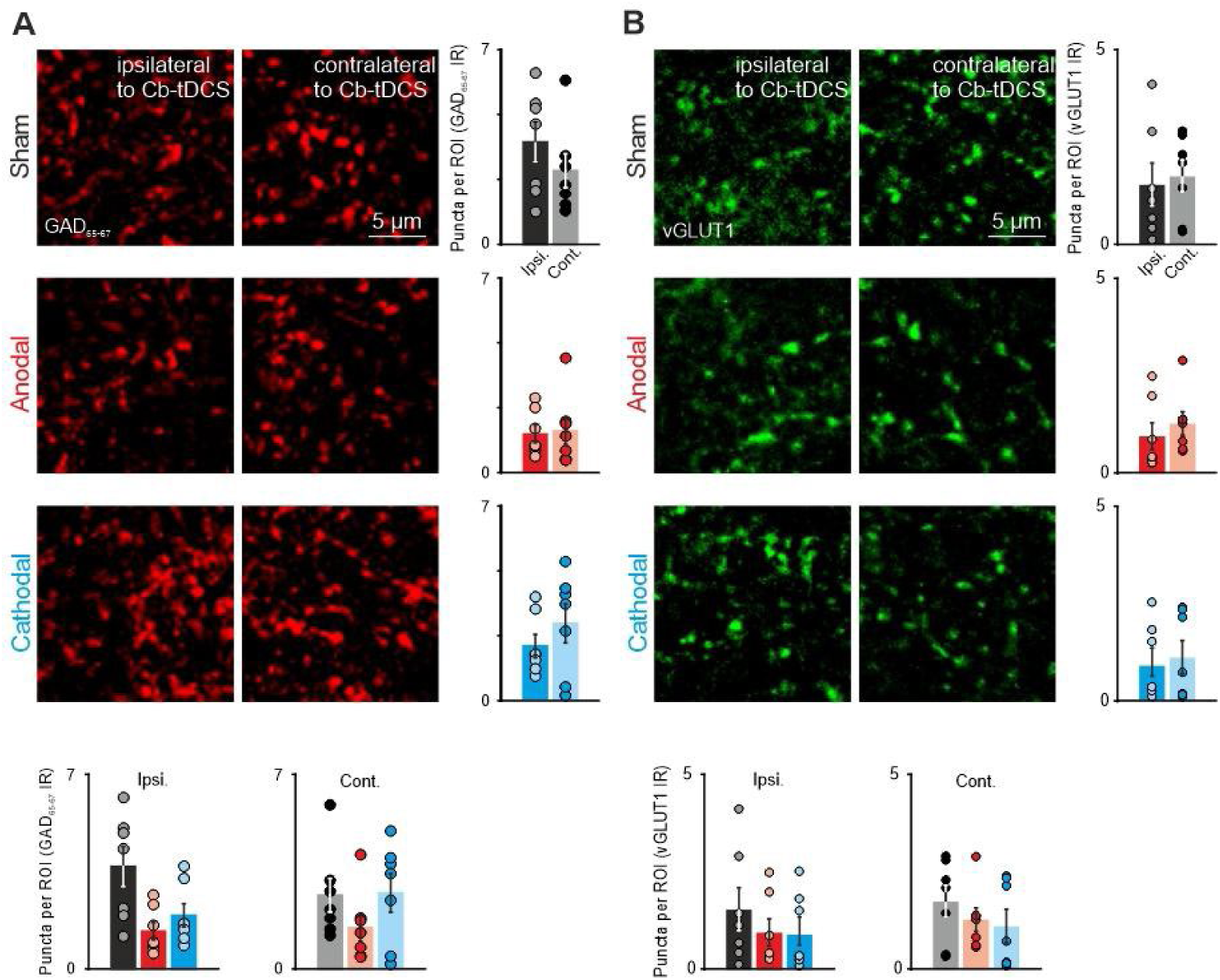
Immunohistochemical immunoreactivity in S1 after 20 min of Cb-tDCS. *A,B*) Representative confocal photomicrographs, quantification, and statistical analysis of GAD65-67 (*A*) and vGLUT1 immunoreactivity (*B*) in the ipsilateral and contralateral S1 hemispheres relative to Cb-tDCS application, after sham stimulation (upper row, N = 7 mice), anodal Cb-tDCS (middle row, N = 7 mice), and cathodal Cb-tDCS (lower row, N = 7 mice). Bottom graphs show the quantification for each hemisphere across stimulation conditions. Error bars represent the standard error of the mean (SEM). Individual data points are overlaid on the bar plots. Statistical analysis was performed using linear mixed-effects models with POLARITY and HEMISPHERE as fixed factors and animal as a random intercept. A significant effect of POLARITY was detected for GAD65-67, whereas no significant effects were found for vGLUT1. GAD65-67: Glutamic acid decarboxylase isoforms 65 and 67; vGLUT1: vesicular glutamate transporter 1; ROI: region of interest.

### Cb-tDCS elicits changes in the oscillatory properties of S1 but does not affect the stimulated region of the cerebellum

Finally, we calculated spectral power ratio values to compare oscillatory activity across animals while accounting for inter-individual variability in baseline power levels. For each animal, power values were averaged and referenced to the corresponding control period. In Crus I/II, spectral power data were analyzed using a linear mixed-effects model with POLARITY (anodal vs cathodal), PERIOD (control vs post-tDCS), and BAND (theta, beta, low-gamma, high-gamma) as fixed factors, and animal as a random intercept. The analysis revealed no significant main effects of POLARITY (F_1,175.776_ = 0.422, p = 0.517), PERIOD (F_1,159.316_ = 0.211, p = 0.646), or BAND (F_3,159.316_ = 0.222, p = 0.881), and no significant interactions among these factors (POLARITY × PERIOD: F_1,159.316_ = 1.899, p = 0.170; POLARITY × BAND: F_3,159.316_ = 0.051, p = 0.985; PERIOD × BAND: _F3,159.316_ = 0.222, p = 0.881; POLARITY × PERIOD × BAND: F_3,159.316_ = 0.051, p = 0.985), indicating that Cb-tDCS did not significantly modify cerebellar spectral power in the analyzed bands (Fig. 8A).

**Figure 8.**
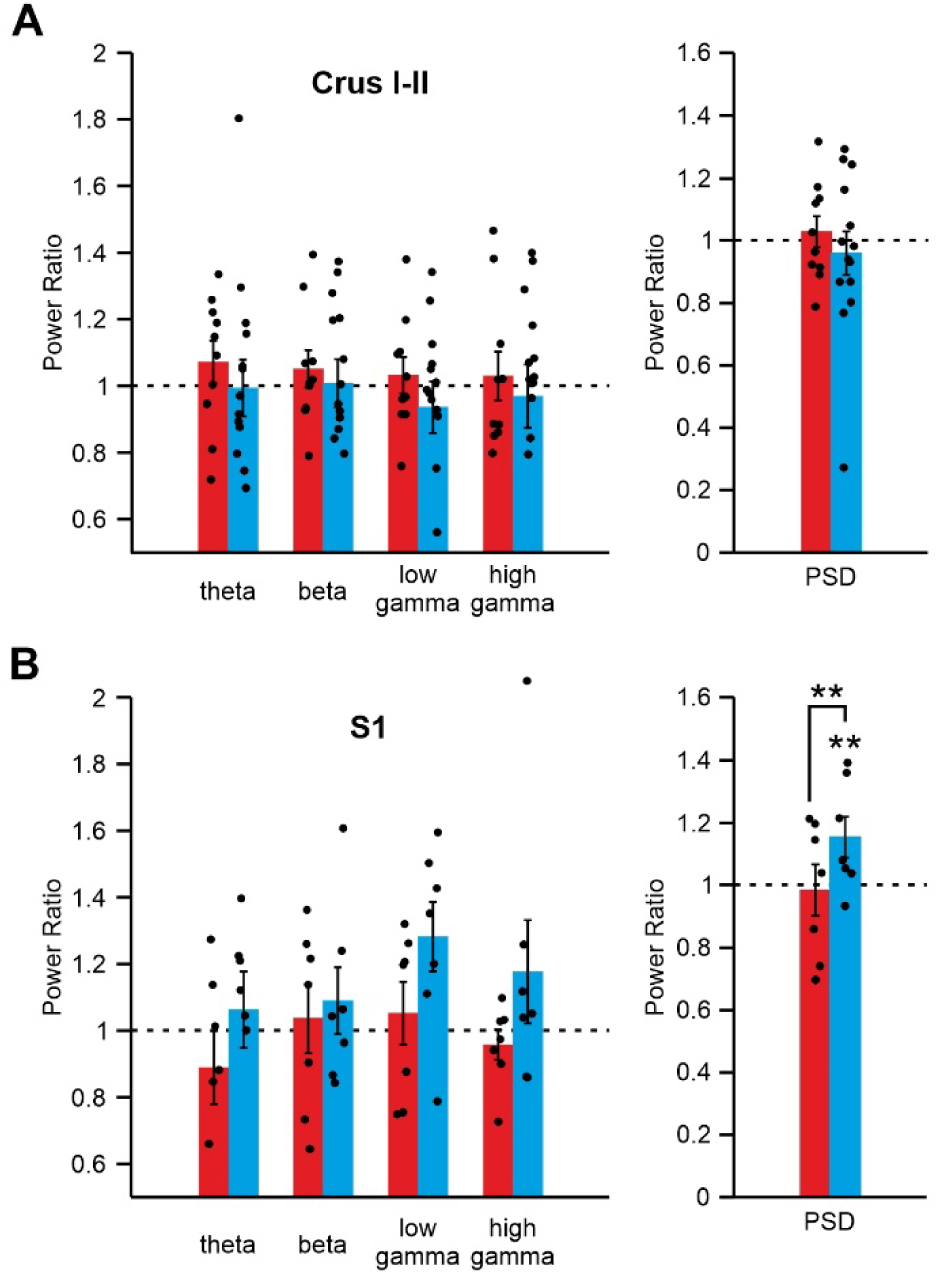
Spectral power analysis of Crus I/II and S1 after Cb-tDCS. Spectral power ratios in the theta, beta, low gamma, and high gamma bands are shown for the Crus I/II (A) and S1 (B) regions, comparing the post-tDCS period (20 min post-tDCS) with the corresponding control period. Red bars indicate anodal Cb-tDCS and blue bars indicate cathodal Cb-tDCS. The dashed black line represents the reference value of the power ratio during the control period. The right panels show the global PSD across the analyzed bands. Statistical analysis was performed using linear mixed-effects models with POLARITY, PERIOD, and BAND as fixed factors, and animal as a random intercept. In Crus I/II, no significant effects were detected (anodal, N = 10; cathodal, N = 14). In S1, spectral power showed a significant effect of POLARITY and a significant POLARITY × PERIOD interaction, driven mainly by increased power after cathodal stimulation in the post-tDCS period (anodal, N = 7; cathodal, N = 7). Asterisks indicate significant differences between anodal and cathodal conditions at the corresponding comparison (**p < 0.01). Error bars represent the standard error of the mean (SEM). Individual data points are overlaid on the bar plots. Theta: 4-10 Hz; beta: 10-30 Hz; low gamma: 30-45 Hz; high gamma: (60-100 Hz).

In S1, the same analysis revealed a significant main effect of POLARITY (F_1,89.075_ = 5.507, p = 0.021) and a significant POLARITY × PERIOD interaction (F1,87.420 = 5.388, p = 0.023), whereas the effects of PERIOD (F_1,87.420_ = 3.526, p = 0.064) and BAND (F_3,87.420_ = 1.153, p = 0.332) were not significant. No significant interactions involving BAND were detected (POLARITY × BAND: F_3,87.420_ = 0.311, p = 0.817; PERIOD × BAND: F_3,87.420_ = 1.153, p = 0.332; POLARITY × PERIOD × BAND: F_3,87.420_ = 0.311, p = 0.817). Bonferroni-corrected pairwise comparisons showed a significant difference between anodal and cathodal stimulation (p=0.021). This difference was not present during the control period (p=0.903), but emerged in post-tDCS (p=0.001). Within-group comparisons further showed that spectral power increased significantly from control to post-tDCS in the cathodal condition (p=0.004), whereas no significant change was detected in the anodal condition (p=0.755) (Fig. 8B). These results indicate that Cb-tDCS did not significantly affect cerebellar spectral power, but induced polarity-dependent post-stimulation changes in S1, driven mainly by an increase after cathodal stimulation. In summary, Cb-tDCS did not significantly modify spectral power in Crus I/II, whereas in S1 it induced polarity-dependent post-stimulation changes, with increased power after cathodal stimulation.

## Discussion

Cb-tDCS has been shown to modulate neural activity not only within the cerebellar region (Manto et al., 2022) but also in cortical areas (Grimaldi et al., 2016). Therefore, the primary objective of this study was to investigate the impact of Cb-tDCS on local cerebellar excitability and on neuronal excitability in S1, the remote cortical site examined here. By understanding how Cb-tDCS influences neuronal responses, we should unravel the intricate functional connectivity of the cerebellum, explore its potential therapeutic applications, and advance our comprehension of brain plasticity. This knowledge holds the key to refining treatment approaches for neurological disorders and fostering the development of innovative interventions that enhance brain function. In this study, we sought to investigate the specific effects of Cb-tDCS on SEPs within distinct cerebellar and cortical regions. By doing so, we aimed to uncover valuable insights into the underlying mechanisms and potential clinical implications of cerebellar stimulation.

### Characterization of cerebellar SEPs

The cerebellum receives sensory information from all parts of the body (Garwicz et al., 1998; Apps & Garwicz, 2005) through mossy fibers (Odeh et al., 2005; Jörntell & Ekerot, 2006) and climbing fibers (Ekerot & Jörntell, 2001). Specifically, peripheral stimulation of the whisker pad evokes a sensory potential in CrusI/II lobules of the cerebellar cortex (Brown & Bower, 2002; Lu et al., 2005; Roggeri et al., 2008) that is characterized by two main components (Fig. 1B): the trigeminal (T) component (Armstrong & Drew, 1980; Bower & Woolston, 1983; Morissette & Bower, 1996; Lu et al., 2005; Roggeri et al., 2008) and the cortical (C) component (Sasaki et al., 1969; Brown & Bower, 2002; Mostofi et al., 2010; Diwakar et al., 2011; Parasuram et al., 2018) inputs. Furthermore, the T component consists of N2-N3 waves that reflect the activity between parallel fibers and PC (Márquez-Ruiz & Cheron, 2012). Following prior research, our study demonstrated that electrical stimulation of the whisker elicited a short-latency SEP in the Crus I/II region, exhibiting comparable latency to the findings reported by Márquez-Ruiz and Cheron in 2012. Since the PC are the sole output of the cerebellar cortex, the amplitude of T component can be used as an index of cerebellar cortex excitability. In our experiments, the intensity of whisker stimulation was adjusted to produce a T component with half of the maximum amplitude, ensuring the observation of potential increases as well as decreases in the evoked potentials during and/or after Cb-tDCS.

### Local effects of Cb-tDCS

In 2009, Galea and colleagues (Galea et al., 2009) demonstrated that the connectivity between the cerebellum and M1 can be increased or decreased, depending on the polarity of tDCS applied to human cerebellum. Since then, Cb-tDCS has been used to modulate motor and non-motor behaviors (Grimaldi et al., 2014, 2016; Oldrati & Schutter, 2018), as well as to understand cerebellar functions (Galea et al., 2011; Ferrucci et al., 2012; Boehringer et al., 2013; Miall et al., 2016) and learning mechanisms (Spampinato & Celnik, 2017). In the initial phase of this study, we examined the local effects of Cb-tDCS on Crus I/II by assessing both immediate and long-term impacts on sensory inputs elicited by electrical stimulation of the whiskers.

Regarding immediate effects, we observed that the amplitude of the T component of the SEP was increased during anodal and decreased during cathodal Cb-tDCS. These results indicate that the immediate effects of Cb-tDCS in Crus I/II are component- and polarity-dependent, with a robust modulation of the trigeminal component and a more limited effect on the cortical component. These findings are consistent with previous *in vitro* experiments conducted in isolated turtle cerebellum (Chan & Nicholson, 1986; Chan et al., 1988) and anesthetized rats (Asan et al., 2020), which demonstrated modulation of PC firing rates when DC currents were applied to the cerebellum. Recent research utilizing high-density neuronal recordings in awake mice further highlighted the significance of PC orientation in the ultimate impact of Cb-tDCS, revealing contrasting firing rate modulation in adjacent PC layers with opposite orientations (Sánchez-León et al., 2025). These results suggest that the final effect of Cb-tDCS on a specific region is highly dependent on the anatomical characteristics of the region of interest. In humans, fMRI studies have yielded mixed results in the context of Cb-tDCS effects. For instance, D’Mello and colleagues (D’Mello et al., 2017) observed that anodal Cb-tDCS increased activation in the right Crus I/II during a complex task involving semantic prediction. Additionally, it enhanced resting-state functional connectivity between hubs associated with the task, specifically the reading and language networks. In contrast, no significant differences in cerebellar cortex activity were reported when Cb-tDCS was applied during a simple motor task like finger tapping (Küper et al., 2019). Interestingly, Küper et al. (2019) found the opposite pattern of modulation in the dentate nuclei after Cb-tDCS. They observed increased activation during cathodal stimulation and a trend toward decreased activation during anodal stimulation. These findings align with the inhibitory effect of cathodal Cb-tDCS on the cerebellar cortex, which results in reduced inhibition of the CN by Purkinje cells, and vice versa. It is important to note that the electric fields generated during Cb-tDCS in humans are typically below 1 V/m (Parazzini et al., 2014; Fiocchi et al., 2016; Rezaee & Dutta, 2019). This contrasts with rodent studies where electric fields can reach up to 60 V/m (Márquez-Ruiz et al., 2014; Sánchez-León et al., 2025). This difference should be taken into account when considering the translational implications of our findings. In humans, computational modeling studies have shown that cerebellar tDCS generates electric fields and current densities that are distributed mainly over the cerebellar cortex, with the resulting intracranial field pattern strongly shaped by head anatomy, electrode montage, and tissue properties (Parazzini et al., 2014; Fiocchi et al., 2016; Rezaee & Dutta, 2019). More broadly, recent consensus recommendations have emphasized that stimulation dose understanding in transcranial brain stimulation should not be reduced to nominal current intensity alone but should instead be understood as a multidimensional construct determined by magnitude, temporal structure, spatial configuration, and the relationship between delivered and received dose (Soleimani et al., 2026). In this context, the higher current densities used in the present mouse preparation were selected to facilitate the detection of acute physiological effects under controlled experimental conditions. Therefore, our results should be interpreted primarily as mechanistic preclinical evidence that cerebellar stimulation can modulate local and downstream sensory processing, rather than as a direct dosimetric model of human cb-tDCS.

As observed in the immediate-effect results in Crus I/II (Fig. 2B), the cortical C component showed more limited modulation than the trigeminal component, although its direction of change was opposite to that of the trigeminal component.. Specifically, anodal Cb-tDCS reduced C component amplitude, whereas cathodal stimulation did not produce a significant change. This observed tendency aligns with our expectations, considering that the cerebellar cortex exerts an inhibitory influence on the cerebral cortex (Batsikadze et al., 2019). Additionally, the C component has been demonstrated to be modulated by activity from S1, as the suppression of S1 activity abolishes the appearance of the C component upon tactile stimulation (Shimuta et al., 2020). Furthermore, diminishing S1 activity lengthens the C component latency, while enhancing S1 activity shortens it (Morissette & Bower, 1996; Brown & Bower, 2002). According to the hypothesis, increasing cerebellar cortex excitability with anodal Cb-tDCS would decrease activity in the CN, which project to S1 via the thalamus (Proville et al., 2014). Consequently, this would lead to a decrease in the excitability of S1, which in turn projects back to the cerebellar cortex through the pontine nuclei (Shinoda et al., 1987; Odeh et al., 2005), resulting in a reduction of cortical C component amplitude (Morissette & Bower, 1996; Brown & Bower, 2002). Conversely, when cathodal Cb-tDCS is applied, it is expected to cause an increase in C component amplitude. Notably, a distinct positive deflection was consistently observed between the T and C components, which was reduced during anodal stimulation and enhanced during cathodal stimulation. While our initial quantification of the C component was performed using a peak-to-peak approach, the presence of this intermediate positivity challenges the interpretation of such measures. This intermediate positivity may reflect rebound excitation of CN following polarity-dependent modulation of PC firing (Person & Raman, 2012). Anodal Cb-tDCS may enhance PC activity and thus increase inhibition of CN, dampening their excitatory rebound and the associated surface positivity. Conversely, cathodal stimulation may reduce PC firing and disinhibit CN, enhancing their output. This raises the possibility that the modulation of this intermediate component—potentially CN-related—may play a key role in shaping the cerebello-cortical dynamics reflected in the SEP. However, given that the mixed-effects analysis revealed a more robust modulation of the trigeminal component than of the cortical component, these mechanistic interpretations should be considered cautiously.

Regarding long-term effects after 20 min of Cb-tDCS, the mixed-effects analysis confirmed that polarity-dependent modulation of the trigeminal component in Crus I/II was prominent during stimulation, but did not provide evidence for robust long-lasting after-effects once stimulation ceased. Consistent with the electrophysiological findings, immunohistochemical analyses did not reveal persistent changes in GAD65-67 or vGLUT1 immunoreactivity in Crus I/II after a single session of Cb-tDCS. During the stimulation period, anodal Cb-tDCS was associated with higher amplitudes than cathodal stimulation, whereas post-stimulation differences were limited and transient. Similar findings have been reported in fMRI studies in humans, where no significant differences in cerebellar cortical activation were observed following Cb-tDCS during a simple motor task (Küper et al., 2019). Importantly, direct electrophysiological evidence assessing persistent local cerebellar changes after a single Cb-tDCS session remains scarce. The lack of significant changes in GAD65–67 and vGLUT1 immunoreactivity in Crus I/II following a single session of Cb-tDCS is also consistent with the idea that cerebellar stimulation effects may be transient, variable, or difficult to detect after stimulation cessation (Grimaldi et al., 2014, 2016). In humans, polarity-dependent modulation of cerebellar-brain inhibition has been demonstrated after cerebellar tDCS, supporting acute effects of stimulation on cerebellar output (Galea et al., 2009). However, evidence for robust and lasting local cerebellar biochemical changes after a single session remains limited. In line with this, magnetic resonance spectroscopy studies in humans did not detect significant group-level changes in cerebellar GABA or glutamate concentrations after a single anodal session, despite observing behavioral effects related to motor memory retention (Jalali et al., 2018). Another study involving a cerebellar-dependent semantic prediction task during Cb-tDCS reported that anodal stimulation led to an increase in activity in CrusI/II, along with enhanced functional connectivity between the task-related hubs (D’Mello et al., 2017). However, human studies utilizing Cb-tDCS often rely on indirect measurements of cerebellar excitability, such as behavioral outcomes or the excitability of connected regions. In these cases, contradictory results are frequently observed (Pope & Miall, 2012; Zuchowski et al., 2014; Miall et al., 2016; Beyer et al., 2017; Batsikadze et al., 2019). There could be several reasons for this disparity, including variations in the orientation of neurons in different cerebellar areas relative to the applied electric field (Grimaldi et al., 2016; Sánchez-León et al., 2023). Furthermore, cerebellar modulation might distinctly affect distantly interconnected regions (Stagg et al., 2018; Miterko et al., 2019). Additionally, the present results underscore the importance of assessing cerebellar responses during the administration of Cb-tDCS, as local cerebellar effects were most evident online and did not persist robustly after stimulation offset. Even with significantly higher electric fields employed in our experiments, no long-term effects were observed in the cerebellar cortex using SEP recordings. Current evidence suggests that anodal cerebellar tDCS may influence the online component of motor skill learning in some paradigms, although the relative contribution of online and offline effects appears to depend on the task and stimulation protocol (Galea et al., 2011; Cantarero et al., 2015; Ehsani et al., 2016; Samaei et al., 2017). In addition, the cerebellum appears to play a particularly relevant role during early phases of motor skill learning and in visuomotor adaptation tasks (Galea et al., 2011; Spampinato & Celnik, 2017). Consistent with the electrophysiological findings, our immunohistochemical analysis of GAD65-67 and vGLUT1 immunoreactivity levels in Crus I/II did not reveal detectable changes following 20 min of Cb-tDCS. This lack of alteration in neurotransmitter levels after Cb-tDCS has also been observed in human experiments utilizing magnetic resonance spectroscopy (MRS) to measure GABA and glutamate levels (Jalali et al., 2018). However, they also did not find any differences during Cb-tDCS, suggesting that the immediate effects of tDCS on the cerebellar cortex might be confined to the polarization induced by the electric field and not correlated with changes in neurotransmitter levels.

### Distant effects of Cb-tDCS

S1 exhibits a high degree of interconnectivity with the cerebellum, as demonstrated by previous studies (Schmahmann, 2001; Ramnani, 2006; Buckner et al., 2011). The extensive characterization of these projections has been achieved through both anatomical investigations (Leergaard et al., 2000) and functional studies (Allen et al., 1979; Shinoda et al., 1987; Swenson et al., 1989; Morissette & Bower, 1996; Brown & Bower, 2002; Nagao, 2004; Odeh et al., 2005; Watson et al., 2009). However, it is only recently that the details and significance of this connection have begun to be elucidated (Proville et al., 2014; Caligiore et al., 2017; Bostan & Strick, 2018). In the present study, the immediate effects of Cb-tDCS on S1 excitability showed a decrease in N1 amplitude during anodal stimulation and an increase during cathodal stimulation, thus following a pattern opposite to that observed in Crus I/II. Moreover, these polarity-dependent effects became more pronounced at the higher stimulation intensity, indicating that the magnitude of remote cortical modulation depends not only on stimulation polarity but also on current strength. No reliable changes in SEP latency were detected, suggesting that the immediate effect of Cb-tDCS in S1 was mainly expressed in response magnitude rather than timing. In line with these findings, a previous animal study demonstrated that manipulating the activity of the posterior cerebellar cortex (Crus I) using chemogenetic techniques resulted in a decrease or increase in the firing rate of neurons in the contralateral parietal association cortex, depending on whether PC activity was increased or decreased, respectively (Stoodley et al., 2017). Furthermore, human studies utilizing indirect measurements have shown that cathodal Cb-tDCS leads to a decrease in the inhibitory influence exerted by the cerebellum on the primary motor cortex (M1), whereas anodal Cb-tDCS has been found to increase this inhibition (Galea et al., 2009; Batsikadze et al., 2019).

Consistent with the immediate results in S1, cathodal Cb-tDCS increased SEP amplitude during stimulation, whereas anodal stimulation reduced it. However, these polarity-dependent effects evolved over time, and the clearest differences between conditions remained concentrated during the stimulation period. After stimulation offset, the temporal profile of S1 responses differed according to polarity, with a limited post-stimulation divergence between anodal and cathodal conditions. One possible interpretation is that homeostatic mechanisms may contribute to shaping these delayed changes in cortical excitability. In line with this interpretation, Proville and colleagues (Proville et al., 2014) showed that optogenetic stimulation of Crus I induced an initial inhibition followed by a rebound activation of the cerebello-thalamo-cortical pathway, resulting in activation of the primary motor cortex (M1). Notably, a comparable delayed pattern was not evident in our recordings from Crus I/II during Cb-tDCS (Fig. 3B). This suggests that the mechanisms underlying these temporal dynamics may involve downstream regions of the cerebello-thalamo-cortical pathway, such as the thalamus or S1 itself. The immunohistochemical analysis conducted in S1 revealed a significant effect of stimulation polarity on GAD65-67 immunoreactivity, with lower values in the anodal group compared with sham. This finding is consistent with the delayed changes in S1 excitability observed after anodal stimulation and suggests that a reduction in inhibitory tone may contribute to the remote cortical effects induced by cerebellar stimulation. Notably, this effect was not significantly modulated by hemisphere, suggesting that the impact of cerebellar stimulation on inhibitory markers in S1 was not restricted to one side under the present experimental conditions. By contrast, vGLUT1 immunoreactivity was not significantly affected, suggesting that the histological effects detected in S1 were more evident for inhibitory than for excitatory markers. The selective reduction in GAD65-67 observed after anodal, but not cathodal, Cb-tDCS may reflect delayed or homeostatic processes associated with post-stimulation cortical adaptation, rather than the immediate modulation observed during stimulation. This interpretation should be made cautiously, given the important methodological differences between direct cortical stimulation and the remote cortical effects induced here by cerebellar stimulation, as well as between MRS-based estimates of GABA concentration and immunohistochemical measures of GAD65-67 expression (Stagg et al., 2009; Jalali et al., 2018). In this context, polarity-specific reductions in cortical GABA after anodal, but not cathodal, tDCS have also been reported in humans (Kim et al., 2014), supporting the possibility that inhibitory markers do not necessarily show identical modulation patterns across polarities. A decrease in GABA after the administration of anodal tDCS over the same area of analysis has been proven in humans (Nandi et al., 2022) and animal (Zhao et al., 2020) studies, but to the extent of our knowledge this is the first evidence that the cerebellar region targeted here can not only modulate the excitability but also neurotransmitter-related markers in S1, an interconnected cortical region.

Taken together, these findings support the presence of distal effects of Cb-tDCS. The results suggest that the effects of non-invasive neuromodulation are not confined to the stimulated region, but can also be detected in S1, which is consistent with an influence on broader interconnected networks (Stagg et al., 2018). This has important implications for both current (Pope & Miall, 2012; Yosephi et al., 2018; Marron et al., 2019) and future (Stoodley et al., 2017; Menardy et al., 2019) therapies that aim to modulate cerebellar networks. These findings highlight the potential utility of targeting the cerebellum for neuromodulatory interventions, as its effects can influence interconnected cortical regions such as S1.

### Impact of Cb-tDCS on local cerebellar and remote S1 oscillatory activity

Oscillatory activity is a fundamental feature of neural processing and reflects coordinated population dynamics across local and distributed circuits, playing a central role in perception and cognition (Engel & Singer, 2001; Tallon-Baudry & Bertrand, 1999; Herrmann et al., 2004; Kahana, 2006). In this context, changes in spectral power may provide complementary information to evoked responses when assessing the impact of neuromodulation on brain networks. In the present study, spectral power analysis revealed no significant changes in Crus I/II, whereas S1 showed a polarity-dependent post-stimulation modulation. Specifically, spectral power in S1 was higher after cathodal than after anodal Cb-tDCS during the post-stimulation period, and the cathodal condition showed a significant increase relative to control. By contrast, no significant post-stimulation spectral changes were detected in the cerebellar region. This pattern is consistent with the SEP results, which also pointed to a stronger expression of delayed effects in S1 than in Crus I/II. Together, these findings suggest that cerebellar stimulation may have a limited long-lasting impact on local oscillatory activity while exerting more persistent consequences on interconnected cortical circuits, in line with the distributed network effects previously described for non-invasive brain stimulation (Stagg et al., 2018). Importantly, evoked responses and spectral power likely capture partially distinct aspects of network function; therefore, the absence of local spectral changes in Crus I/II does not contradict the presence of immediate SEP modulation in the same region. Rather, it suggests that Cb-tDCS may influence local sensory responsiveness without necessarily inducing sustained changes in ongoing oscillatory power. Importantly, the spectral effects observed in S1 were not restricted to a specific frequency band in the present analysis, suggesting a broader post-stimulation modulation of cortical network state rather than a selective effect confined to a single oscillatory range. The divergence between local and remote effects also suggests that Cb-tDCS may engage fast mechanisms at the site of stimulation and slower network-level processes in downstream cortical targets. From a mechanistic perspective, these remote oscillatory effects are compatible with the view that cerebellar stimulation can influence cortical processing through distributed cerebello-thalamo-cortical pathways (Galea et al., 2009; Proville et al., 2014), thereby reshaping post-stimulation network dynamics outside the stimulated region itself. In addition, our previous work using direct tDCS over S1 in awake mice showed polarity-dependent effects on cortical oscillatory activity (Sánchez-León et al., 2021), suggesting that remote modulation induced by cerebellar stimulation does not necessarily reproduce the same spectral signature observed after direct cortical stimulation. This interpretation is also consistent with previous evidence showing that cerebellar stimulation can modulate activity in distributed cortical networks beyond the stimulated region itself (Stagg et al., 2018; D’Mello et al., 2017). Overall, these findings indicate that the oscillatory consequences of Cb-tDCS are expressed more prominently in remote cortical circuits than in the stimulated cerebellar cortex, highlighting the importance of considering distributed network effects when interpreting cerebellar neuromodulation.

In conclusion, the study provides compelling evidence that Cb-tDCS effectively modulates the amplitude of SEPs in both the cerebellum and S1. Notably, immediate opposite effects were observed during the application of Cb-tDCS, suggesting a bidirectional influence. On the other hand, long-term effects were found to extend primarily in S1, indicating that Cb-tDCS can exert sustained effects beyond the stimulated cerebellar region. These findings significantly advance our understanding of the neural mechanisms underlying Cb-tDCS and its potential for modulating cerebellar-brain interactions.

## Data availability

The datasets generated and analyzed during the current study are publicly available in Zenodo at https://doi.org/10.5281/zenodo.20306373.

## Author contributions

C.A.S-L and J.M-R conceived the original idea and designed the experiments. C.A.S-L, I.C, A.J-D and J.M-R performed the experiments and the data analysis. C.A.S-L and J.M-R wrote the paper. G.C and J.F.M assisted in the experimental design and in the interpretation of the results. All the authors contributed to the final edition of the manuscript.

## Declaration of generative AI and AI-assisted technologies in the writing process

During the preparation of this work, the authors used ChatGPT-4o to enhance language clarity and polish the writing. After using this tool, the authors reviewed and edited the content as needed and take full responsibility for the content of the publication.

